# Distinct roles of monkey OFC-subcortical pathways in adaptive behavior

**DOI:** 10.1101/2023.11.17.567492

**Authors:** Kei Oyama, Kei Majima, Yuji Nagai, Yukiko Hori, Toshiyuki Hirabayashi, Mark A G Eldridge, Koki Mimura, Naohisa Miyakawa, Atsushi Fujimoto, Yuki Hori, Haruhiko Iwaoki, Ken-ichi Inoue, Richard C Saunders, Masahiko Takada, Noriaki Yahata, Makoto Higuchi, Barry J Richmond, Takafumi Minamimoto

## Abstract

To be the most successful, primates must adapt to changing environments and optimize their behavior by making the most beneficial choices. At the core of adaptive behavior is the orbitofrontal cortex (OFC) of the brain, which updates choice value through direct experience or knowledge-based inference. Here, we identify distinct neural circuitry underlying these two separate abilities. We designed two behavioral tasks in which macaque monkeys updated the values of certain items, either by directly experiencing changes in stimulus-reward associations, or by inferring the value of unexperienced items based on the task’s rules. Chemogenetic silencing of bilateral OFC combined with mathematical model-fitting analysis revealed that monkey OFC is involved in updating item value based on both experience and inference. In vivo imaging of chemogenetic receptors by positron emission tomography allowed us to map projections from the OFC to the rostromedial caudate nucleus (rmCD) and the medial part of the mediodorsal thalamus (MDm). Chemogenetic silencing of the OFC-rmCD pathway impaired experience-based value updating, while silencing the OFC-MDm pathway impaired inference-based value updating. Our results thus demonstrate a dissociable contribution of distinct OFC projections to different behavioral strategies, and provide new insights into the neural basis of value-based adaptive decision-making in primates.

## Introduction

To survive in a constantly changing world, animals naturally adapt quickly to new environments and adjust their behavior to maximize the benefits. This involves making decisions that will lead to maximum subjective benefit, based on the changing relationships between specific events and outcomes, and thus requires knowing the current worth of each option when making a choice. Typically, an item’s worth is learned through direct experience, a process often explained through the concept of classical reinforcement learning(1). However, animals with highly developed brains, particularly primates, have also evolved the ability to infer the value of unexperienced events/items from their knowledge of the world. This ability is described by the rule or theory of shifting relationships. For example, if a monkey is eating a banana and notices that it is ripe, it may be able to infer that other nearby bananas are also ripe, based on its knowledge of how fruits ripen. Optimal decision-making relies on a balance between experience- and inference-based behavioral strategies. The primate orbitofrontal cortex (OFC) is thought to contribute to such adaptive behavior by leveraging both direct experience and knowledge of the current context. The OFC has long been thought to play an essential role in encoding the subjective values of alternative events/items that guide subsequent decision-making, and in learning/updating these values by integrating past experiences as consequences of our choices(2-6). At the same time, the OFC has also been shown to be necessary for inferring value based on mental simulation of outcomes, even in the absence of direct experience, by generalizing knowledge of the current situation or environment(7-9). A recent report suggests that the OFC regulates the balance between these two valuation strategies, rather than simply initiating one or the other(10). Thus, there is ongoing debate about the core function of the OFC in adaptive behavior.

The complexity of OFC function might arise from its interactions with other brain regions through direct anatomical connections(11). For example, subcortical structures, such as the rostromedial part of the caudate nucleus (rmCD) and the medial part of the mediodorsal thalamus (MDm), receive direct projections from the OFC(12-14). Lesions to these areas have been shown to produce deficits that are similar, but not identical, to those produced by OFC lesions(15-20) with a tendency that rmCD and MDm are particularly involved in value-updating based on experience and inference, respectively, suggesting that they have overlapping yet distinct roles in adaptive behavior. These findings have raised the possibility that two pathways originating from OFC, namely the OFC-rmCD and OFC-MDm pathways, are needed for different value-updating strategies. To investigate this possibility, it is essential to independently manipulate the prefronto-subcortical circuits, which is technically challenging, especially in behaving nonhuman primates.

To investigate the causal roles of the OFC and its originating pathways in these two types of valuation, we use a chemogenetic tool called designer receptors exclusively activated by designer drugs (DREADDs)(21). This tool allows neurons to be silenced by activating an inhibitory DREADD (hM4Di) following systemic administration of a DREADD agonist. Additionally, local agonist infusion that activates hM4Di expressed at axon terminals can suppress synaptic transmission(22, 23). By combining these techniques with positron emission tomography (PET) as an in vivo imaging tool for hM4Di-positive projection sites, we have previously developed imaging-guided chemogenetic synaptic-silencing that is dramatically more efficient and accurate, especially when applied to nonhuman primates(24). Leveraging this technique and a model-fitting analysis in a reinforcement learning framework, the present study addresses the contributions of these two OFC-subcortical pathways to different value-updating strategies. Our results suggest that experience- and inference-based strategies for updating stimulus-reward associations rely on the OFC-rmCD and OFC-MDm pathways, respectively.

## Results

### Experience- and inference-based behavioral adaptation in multi-reward value-updating tasks

To address the question of how the OFC and its projections to subcortical structures contribute to behavioral adaptation through experience- and inference-based valuation, we devised two behavioral tasks for macaque monkeys: NOVEL and FAMILIAR tasks, respectively (Fig. 1A). In both tasks, the monkeys were required to choose either of two presented visual stimuli (out of a set of five abstract images), each of which was associated with 1, 2, 3, 4, or 5 drops of juice. The order of associations was reversed within a session (Fig. 1A). To maximize their reward, monkeys had to learn the values of the visual stimuli and then update them following subsequent changes in stimulus-reward associations. In the NOVEL task, which was aimed to assess experience-based updating, a new set of stimuli was introduced in each session, thus requiring the monkeys to learn new stimulus-reward associations as well as the association reversal that was imposed mid-session (90 trials after the beginning of each session). After several months of training, two monkeys (Mks #1 and 2) were able to learn new stimulus-reward associations within 80-90 trials and adapted to their reversal within 30-50 trials (Fig. 1B, right). In the FAMILIAR task, which was aimed to assess inference-based updating, a fixed set of five visual stimuli was used throughout the experiments, and the reversals were imposed several times after performance reached a predetermined criterion (see Methods). Following several months of training on this task, the monkeys were able to adapt to the reversal within 3-5 trials (Fig. 1C, right). They even showed optimal choice for “unexperienced” stimulus-reward associations after experiencing the other associations following the reversal (for details, see latter section), indicating that they solved this task based on inference, that is, their prior knowledge of the limited patterns of stimulus-reward associations.

**Fig. 1.**
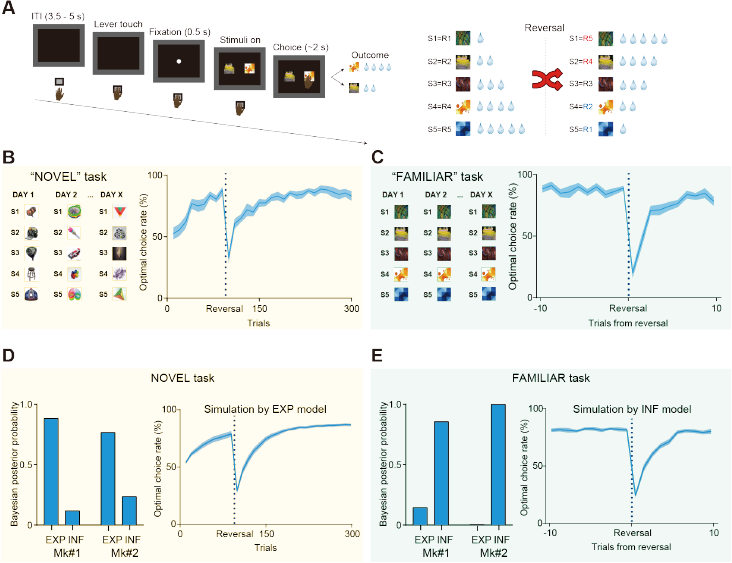
Experience- and inference-based value updating in multi-reward reversal learning tasks. (A) Sequence of a trial (left) and the reversal rule for stimulus-reward associations (right). S1-S5 represent the identity of each stimulus and R1-R5 represent the amount of reward (1 to 5 drops of juice) associated with each stimulus. (B and C) Examples of stimulus sets and baseline performance for the “NOVEL” (B) and “FAMILIAR” (C) tasks. Data were averaged for two monkeys (N = 16 and 14 sessions; 8 and 7 sessions for each monkey in NOVEL and FAMILIAR tasks, respectively). Solid lines and shaded area represent the mean and s.e.m, respectively. Data for the FAMILIAR task were truncated to show those around the reversals and were averaged across reversals. (D and E) Bayesian posterior probability calculated for “EXP” and “INF” models given the behavioral data in each task (left) and the behavioral simulation by the model with higher posterior probability (right); EXP model for the NOVEL task (D) and INF model for FAMILIAR task (E), respectively.

To examine whether the monkeys solved these tasks based on experience- or inference-based strategies, we conducted simulations using two types of reinforcement learning models (see Methods for details); one assuming that the values were updated through direct experience based on standard model-free reinforcement learning that was driven by reward prediction errors (“EXP” model), and one assuming that the monkeys had a priori knowledge of the two possible stimulus-reward association patterns (1,2,3,4,5 or 5,4,3,2,1 drops) through daily training, and that prediction errors drove the transition between these two already learned value sets, allowing them to infer the values of any unexperienced stimuli (“INF” model). As expected, this analysis revealed that behavior during the NOVEL task was better explained by the EXP model (Fig. 1D), while that during the FAMILIAR task was better explained by the INF model (Fig. 1E). These results suggest that monkeys “solved” the two tasks with different strategies — experience-based for the NOVEL task and inference-based for the FAMILIAR task.

### OFC silencing impairs both experience- and inference-based strategies

Next, we chemogenetically inactivated the OFC to determine whether it contributes to experience- and/or inference-based valuation strategies. First, we introduced the inhibitory DREADD, hM4Di, bilaterally into the lateral OFC (Brod-mann’s area 11/13) of the two monkeys via injections of an adeno-associated virus (AAV) vector (AAV2-CMV-hM4Di and AAV2.1-CaMKII-hM4Di-IRES-AcGFP for Mks #1 and 2, respectively) (Fig. 2A). Several weeks after the injections, we non-invasively visualized hM4Di expression using PET imaging with DREADD-PET tracers. In both monkeys, we consistently observed increased PET signal in the bilateral OFC (Fig. 2A; fig. S1A), which was confirmed by post-mortem immunohistochemistry to reflect hM4Di expression. We then examined the effect that silencing the OFC had on behavior; task performances were compared after systemically administering either a vehicle control or the DREADD agonist deschloroclozapine (DCZ) (Fig. 2B). While silencing the OFC did not alter performance on the NOVEL task before reversal of the stimulus-reward contingencies (acquisition phase), it did impair performance following the reversal. This was particularly true in the early phase (Fig. 2C) and was consistently observed in both monkeys (fig. S2A,B). Similarly, OFC silencing also impaired performance on the FAMILIAR task just after reversals (Fig. 2D, fig. S2C,D). To quantify the silencing effect in both tasks, we fitted two behavioral models to the behavioral data following DCZ and vehicle administration. Model-fitting analysis revealed that the impaired performance after OFC silencing could be attributed to a decrease in the learning rate (α) after the reversals (Fig. 2E,F, see Methods for details), but not to the change in the extent of exploration (the inverse temperature, β; fig. S3A,D), suggesting that silencing the OFC led to deficits in value-updating when using either strategy.

**Fig. 2.**
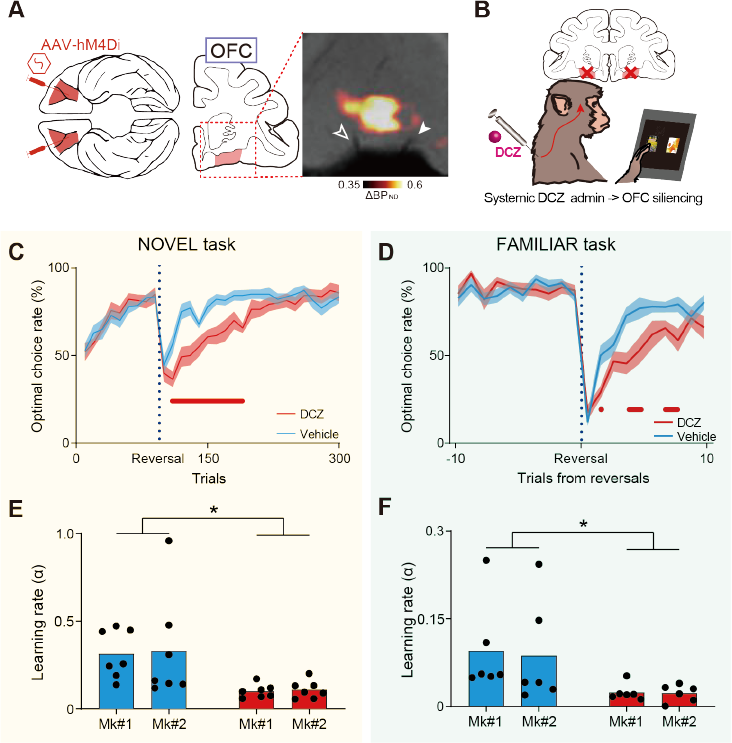
Chemogenetic silencing of the bilateral OFC impaired experience- and inference-based value updating. (A) Injection area (bilateral OFC, Brodmann’s area 11/13) and PET image showing hM4Di expression in Mk2. (B) Shema of OFC silencing per se. DCZ (100 ug/kg) was systemically injected intramuscularly. (C and D) Behavioral effects of chemogenetic OFC silencing on NOVEL (C) and FAMILIAR (D) task performance. Data for vehicle injections (cyan) and DCZ injections (red) are shown. The optimal choice rate (n = 7 and 6 sessions for each treatment in each monkey for the NOVEL and FAMILIAR tasks, respectively) following DCZ (red) and vehicle injections (cyan) was averaged across two monkeys. The red marks below each graph indicate trial numbers after reversal for which significant differences between vehicle and DCZ conditions were observed (t-test, p < 0.05). (E and F) Estimated learning rates in the NOVEL task with the EXP model (E, two-way ANOVA, treatment × monkey, main effect of treatment, F(1,24) = 11.3, p = 2.6 × 10-3; subject, F(1,24) = 0.04, p = 0.84; interaction, F(1,24) = 0.004, p = 0.95)) and in the FAMILIAR task with the INF model (F, treatment, F(1,20) = 7.5, p = 1.3 × 10-2; subject, F(1,20) = 0.04, p = 0.85; interaction, F(1,20) = 0.02, p = 0.90). Asterisks: p < 0.05 for significant main effect of treatment.

We conducted four additional experiments to confirm that the observed behavioral changes were due to the loss of normal OFC function that is involved in adapting to the shift in the task context, and which requires updating the values of the external stimuli. First, to confirm that healthy OFC function is required at the time of the reversal, but not when learning the stimulus-reward associations or during any other process in the acquisition phase, we administered DCZ after the acquisition phase (just before reversal, see Methods for details). This manipulation produced deficits similar to those observed when DCZ was administered before the beginning of each session (fig. S4A), suggesting that the effects of OFC silencing were limited to the reversal of stimulus-reward associations. Second, we confirmed that DCZ alone did not significantly affect behavioral performance before the introduction of hM4Di (Mk #2, fig. S4B), indicating that the effects observed after DCZ administration were due to DREADD activation. Third, we confirmed that OFC silencing did not significantly impact simple reversal learning (fig. S5), consistent with a previous lesion study(25). Fourth, to test whether the OFC is also essential for situations in which knowledge-based value-updating is required and the cognitive load is high, but the item values are binary (i.e., reward or no reward), we examined the effects of OFC silencing using an analog of the Wisconsin Card Sorting Test(26) (Mk #2, fig. S6B). In this case, we found that silencing the OFC had no effect on task performance (fig. S6C), suggesting that it is required specifically when updating external items with complex, non-binary values. Importantly, chemogenetic silencing of the OFC remained effective at the end of all experiments, including these control experiments and the pathway-selective manipulations (see below) (fig. S6A), as demonstrated by the significant effects of DCZ administration on performance during the final devaluation task (fig. S6D,E), which is one of the most common tasks requiring normal OFC function(7, 27). Taken together, these results suggest that the OFC is essential for adapting behavioral responses that are specifically contingent upon being able to update multiple values of multiple external stimuli.

### The OFC-rmCD and OFC-MDm pathways are necessary for experience- and inference-based value-updating, respectively

Having demonstrated that the OFC is essential for both experience- and inference-based behavioral adaptation, the next question is whether these different strategies are governed by separate neural pathways. To answer this, we conducted a chemogenetic pathway-selective manipulation wherein information flow can be temporarily inactivated by local agonist infusion into axonal terminals expressing hM4Di. It is generally challenging to precisely localize and target axonal projection sites in vivo in monkeys, which have relatively large and complexly shaped brains. We overcame this obstacle by using an imaging-guided chemogenetic technique(24) in which PET allows localization of the projection sites for hM4Di-positive OFC neurons. Aside from the PFC, subtraction PET images (post-AAV injection minus pre-AAV injection) showed increased PET signals in the striatum and the thalamus, specifically in the rmCD, MDm, and the medial part of the putamen (Fig. 3A,B; fig. S1B-D). These regions colocalized with GFP-positive axon terminals under immunohistological examination (fig. S1B-D). In contrast, the hM4Di signal was not clearly observed via PET or histology in other brain regions which are known to also receive projections from the OFC, such as the amygdala, and was not comparable to what we observed in these three regions (fig. S1E). Thus, our next experiments focused on the projections from the OFC to the three DREADD-positive terminal regions.

**Fig. 3.**
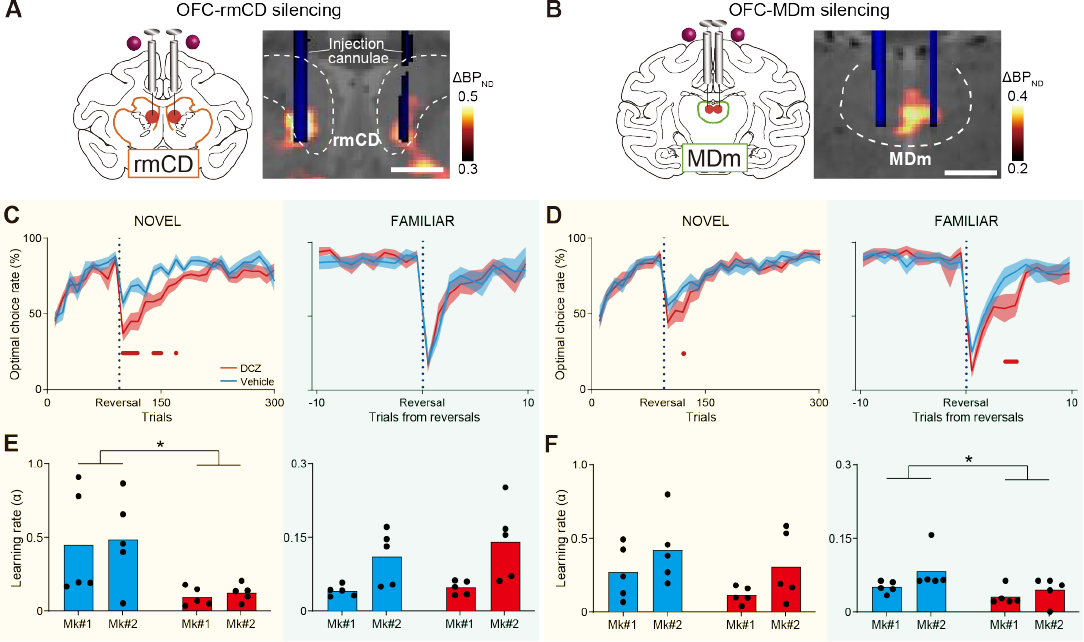
OFC-rmCD and OFC-MDm pathways are necessary for experience- and inference-based value-updating, respectively. (A and B) vChemoge-netic silencing of the OFC-rmCD (A) and OFC-MDm (B) pathways by local DCZ infusion into either bi-lateral rmCD or MDm, specifically at hM4Di-positive OFC terminal sites. A CT image showing the infusion cannulae (blue) overlaying a structural MR image (grey), and a PET image showing a high [11C]DCZ binding region (hM4Di expression, hot color) ob-tained from Mk2. The dashed lines represent the borders of the caudate nucleus and mediodorsal tha-lamus, respectively. (C and D) Optimal choice rate (n = 10 sessions for each treatment, 5 sessions in each monkey) in the NOVEL (left) and FAMILIAR (right) tasks after silencing the OFC-rmCD (C) and OFC-MDm (D) pathways. Conventions are the same as in Fig. 2. (E and F) Estimated learning rates after silencing the OFC-rmCD pathway during the NOVEL (E, left, treatment, F(1,16) = 11.0, p = 4.4 × 10-3; subject, F(1,16) = 0.10, p = 0.76; interaction, F(1,16) = 0.0022, p = 0.96) and FAMILIAR (E, right, treat-ment, F(1,16) = 0.74, p = 0.40; subject, F(1,16) = 14.1, p = 1.8 × 10-3; interaction, F(1,16) = 0.30, p = 0.59) tasks, and the OFC-MDm pathway during the NOVEL (F, left, treatment, F(1,16) = 2.5, p = 0.14; subject, F(1,16) = 3.9, p = 0.07; interaction, F(1,16) = 0.059, p = 0.81) and FAMILIAR (F, right, treatment, F(1,16) = 5.7, p = 2.9 × 10-2; subject, F(1,16) = 3.7, p = 0.07; interaction, F(1,16) = 0.61, p = 0.45) tasks. Scale bars: 5 mm.

To reversibly block neuronal transmission from the OFC, we infused DCZ into one of the three terminal regions under the guidance of MR, PET, and CT imaging (Fig. 3A,B). Compared with the control vehicle infusion, local DCZ infusion into the rmCD impaired performance on the NOVEL task just after reversals, similar to the systemic DCZ injections (Fig. 3C, left). However, this was not the case for the FAMILIAR task (Fig. 3C, right). Conversely, DCZ infusion into MDm resulted in relatively minor impairment on the NOVEL task (Fig. 3D, left), but significant impairment on the FAMILIAR task (Fig. 3D, right). Although we also injected DCZ into the medial part of the putamen, where we found weak PET and histological signals (fig. S1D), this did not affect performance on the NOVEL task (fig. S7). We therefore focused further tests on the OFC to rmCD and OFC to MDm pathways. Model fitting analysis using two reinforcement learning models revealed that silencing the OFC-rmCD pathway significantly reduced the learning rate during the NOVEL task (EXP model; Fig. 3E, left), but not during the FAMILIAR task (INF model; Fig. 3E, right). Conversely, silencing the OFC-MDm pathway had no impact on the learning rate during the NOVEL task (Exp model; Fig. 3F, left), but significantly reduced it during the FAMILIAR task (INF model; Fig. 3F, right). Similar to OFC silencing, silencing either pathway did not affect the inverse temperature (Extended Data Fig. 3b-f).

Damage to the OFC in humans has been associated with increased impulsivity(28). In monkeys, OFC inactivation or lesioning has resulted in faster reaction times on experimental tasks(29), which is generally interpreted as a sign of impulsivity or lack of control. To assess whether silencing the OFC and its projections affected impulsivity in our task context, and if so, whether this behavioral change is related to the observed impairment in performance, we examined reaction times after each type of silencing (fig. S8). OFC silencing resulted in shorter reaction times for both monkeys during all task phases, but we did not observe any direct relationship between reaction time and performance (Extended Data Fig. 8a). In contrast, silencing the OFC-rmCD and OFC-MDm pathways induced complex and contradicting results; silencing the OFC-rmCD pathway increased reaction time (fig. S8B), whereas silencing the OFC-MDm pathway did not influence reaction time (fig. S8C). Although these effects on reaction time differed, we can conclude that the difficulty in updating item values was not related to increased impulsivity.

Taken together, selective silencing of the two OFC projections indicated that both are involved in updating stimulus values; the OFC-rmCD pathway is needed when updating based on direct experience of stimulus-reward associations, whereas the OFC-MDm pathway is needed when updating based on inference from previously learned knowledge.

### The OFC-rmCD and OFC-MDm pathways differentially contribute to the sensitivity to past outcomes

Our results suggest that the two pathways contribute distinctly to different behavioral strategies. To further corroborate this dissociation, we perform a learning-model agnostic analysis in which we decompose trials into specific events related to each behavioral strategy. First, we focused on experience-based updating during the NOVEL task. If the ability to update values based on past experience is impaired, the behavior following unpredicted positive experiences (i.e., obtaining a good result after choosing the previously unchosen option) should favor repeating the same choice, and vice versa following negative experiences (Fig. 4A). Consistent with the model-fitting analysis, silencing the OFC or the OFC-rmCD pathway significantly impaired performance following both positive and negative experiences (Fig. 4C,D), suggesting that their contribution to updating stimulus-reward associations is based on both positive and negative experiences. In contrast, silencing the OFC-MDm pathway only impaired performance following negative experiences (Fig. 4E), likely reflecting minor deficits at the immediate post-reversal period (Fig. 3D). Similarly, asymmetric deficit was induced by silencing the OFC-MDm pathway during the FAMILIAR task (fig. S9).

**Fig. 4.**
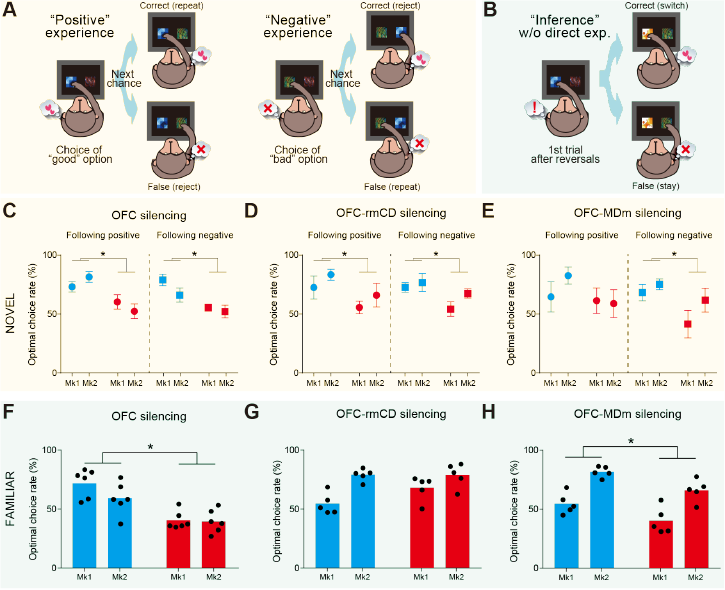
Silencing of OFC-rmCD and OFC-MDm pathways differentially affected the sensitivity to past outcomes. (A and B) Schematic drawings showing the trials following positive (left) and negative (right) experience (A), and the “inference” trial following a single experience of the reversal of stimulus-reward associations (B) for the NOVEL and FAMILIAR tasks, respectively. (C to E) Averaged optimal choice rate for trials following positive (left) and negative (right) outcomes for OFC silencing (C) (positive: treatment, F(1,24) = 14.8, p = 7.9 × 10-4; subject, F(1,24) = 0.001, p = 0.97; interaction, F(1,24) = 2.2, p = 0.15; negative: treatment, F(1,24) = 14.3, p = 9.1 × 10-4; subject, F(1,24) = 2.7, p = 0.11; interaction, F(1,24) = 0.94, p = 0.34), OFC-rmCD silencing (D) (positive: treatment, F(1,16) = 4.8, p = 4.3 × 10-2; subject, F(1,16) = 1.8, p = 0.20; interaction, F(1,16) = 0.001, p = 0.97; negative: treatment, F(1,16) = 6.0, p = 2.6 × 10-2; subject, F(1,16) = 2.4, p = 0.14; interaction, F(1,16) = 0.65, p = 0.43), and OFC-MDm silencing (E) (positive: treatment, F(1,16) = 1.5, p = 0.23; subject, F(1,16) = 0.51, p = 0.48; interaction, F(1,16) = 0.90, p = 0.36; negative: treatment, F(1,16) = 5.2, p = 3.7 × 10-2; subject, F(1,16) = 2.4, p = 0.14; interaction, F(1,16) = 0.56, p = 0.46). (F to H) Averaged optimal choice rate for inference trials after OFC silencing (F) (treatment, F(1,20) = 33.1, p = 1.3 × 10-5; subject, F(1,20) = 2.5, p = 0.13; interaction, F(1,20) = 1.7, p = 0.21), OFC-rmCD silencing (G) (treatment, F(1,16) = 2.7, p = 0.12; subject, F(1,16) = 19.6, p = 4.2 × 10-4; interaction, F(1,16) = 2.9, p = 0.11), and OFC-MDm silencing (H) (treatment, F(1,16) = 14.2, p = 1.7 × 10-3; subject, F(1,16) = 43.7, p = 6.0 × 10-6; interaction, F(1,16) = 0.021, p = 0.88).

Next, we focused on inference-based updating during the FAMILIAR task. We found that the monkeys could adapt to the new stimulus-reward association after a single experience of a reversal (1st trial, Fig. 4B, left), even when the subsequent trial did not include options that appeared in the previous trial (inference trial, Fig. 4B, right). Monkeys exhibited a greater percentage of optimal choices in the inference trial than in the 1st trial (inference vs. first trials; Mk #1: 72.2% vs 19.6%; Mk #2: 59.3% vs 17.7%) under the baseline control conditions, indicating that they inferred the stimulus-reward associations without needing direct experience. This inference-based behavioral adaptation was significantly impaired following silencing of the OFC or the OFCMDm pathway (Fig. 4F,H). In contrast, silencing the OFC-rmCD pathway had no effect on the inference trials (Fig. 4G). Taken together, these results support the conclusion that the OFC-rmCD pathway is essential for updating value via positive and negative experiences, whereas the OFC-MDm pathway is selectively involved in behavioral changes following negative experiences—a capacity that is critical for rapid behavioral adaptation using an inference-based strategy based on prior knowledge of the situation.

## Discussion

Here, by combining chemogenetic silencing of individual neural pathways with a model-fitting approach, we demonstrate that the two natural strategies for updating subjective value of external stimuli rely on two distinct neural pathways, each originating in the OFC and projecting to a different subcortical brain regions. Silencing the OFC-rmCD pathway impaired performance when monkeys updated option values through direct experience of both positive and negative changes of stimulus-reward associations, whereas silencing the OFC-MDm pathway impaired performance when monkeys updated the values based on inference that is guided by negative experience which could notice the change of the task context. This dissociable contribution of neural pathways provides new insights into the neural basis of value-based adaptive decision-making in primates.

The OFC is widely thought to play a central role in updating and maintaining information about possible outcomes(6, 11). Both lesion and recording studies have shown that the OFC is critical for updating associations between stimulus/action and outcome, but not for acquiring such associations(20, 30), which was also the case in our study (Fig. 2). Previous research has also suggested that the OFC is required for reversal learning in various species(31-33). However, recent studies have challenged this view by showing that selective lesion/inactivation of the OFC in macaque monkeys had no effect in a simple reversal learning paradigm(25, 34), a result consistent with our observations (fig. S5). Recently, another study suggested that the OFC is essential for updating the desirability (i.e., quality or quantity) of reward-associated stimuli, but not their availability (i.e., the probability of receiving rewards)(20). Our current results, in which OFC silencing severely impaired performance on a behavioral task that requires monkeys to associate multiple stimuli with multiple reward amounts and to update their associations, support this view. Notably, OFC silencing did not impact performance on the Wisconsin Card Sorting Task (WCST, fig. S6), which is commonly used to assess behavioral flexibility. Although previous lesion studies in monkeys have reported deficits in WCST performance following aspirating lesions(20), subsequent studies have suggested that these deficits may have been due to damaged fibers that pass near the OFC(25). Because outcomes on the WCST are typically binary, either receiving a reward or not, our results suggest that primate OFC, at least the lateral part that we focused on in this study (Brodmann’s area 11/13), is specialized for representing stimulus-reward associations that are based on the desirability of possible outcomes.

Recent research into the role of the OFC has highlighted a specialized role in inference-based, or model-based, learning. This idea has been supported by a number of recording and lesion studies, which have consistently demonstrated the involvement of the OFC in such learning processes(35, 36). However, other recent reports suggest that rodent OFC, or the ventral PFC in humans, is not simply engaged in either experience- or inference-based value-updating strategies, but rather in regulating the switch between them according to current contexts(10, 37). Our present results are consistent with and extend this idea; we can now assign distinct roles for at least two different OFC projections to subcortical structures (rmCD and MDm). To the best of our knowledge, the present study is the first report in primates that has directly investigated the dissociable roles of OFC-originating pathways. Importantly, the two pathways we focused on form part of a broader circuit for value-updating. Therefore comprehensively examining the other circuity, such as the OFC-amygdala pathway(38-40) and the OFC-sensory cortex pathway(41), is essential for a deeper understanding of value-guided behavior.

In the current study, we were able to dissect the functions of the OFC-subcortical circuits by using the imaging-guided pathway-selective synaptic silencing technique that we recently developed(24). Interestingly, in that study, we also observed a functional dissociation in frontal cortex pathways, in that case originating in the dorsolateral prefrontal cortex (dlPFC) and projecting to either the CD or MD thalamus. Although the OFC and dlPFC are located in different parallel cortico-basal ganglia-thalamo-cortical circuits(42), the observed similarity that different subcortical pathways have different functions suggests that the parallel prefrontal-subcortical networks might share a common neural basis for adaptive behavior. Future studies should aim to better understand the overall architecture of these networks by identifying where and how they communicate with each other and how the information processed in each network is integrated.

In summary, leveraging the technical advantage of imaging-guided pathway-selective chemogenetic silencing, we have demonstrated the dissociable contributions of two OFC-subcortical pathways to different value-updating strategies. The identification of causal relationships between a specific neural pathway and a cognitive function, as demonstrated in this study, can compliment what we have learned from human studies, but which cannot be directly tested in humans for ethical reasons. Thus, in addition to providing new insights into the neural basis of value-based adaptive decision-making in nonhuman primates, our results have implications for psychiatric conditions associated with malfunctioning prefrontal-subcortical networks, such as obsessive-compulsive disorder.

## ACKNOWLEDGEMENTS

Two Japanese monkeys were provided by the Japan MEXT National Bio-Resource Project “Japanese Monkeys”. We thank Drs. Brian Russ and Junya Hirokawa for their valuable comments on an earlier version of the manuscript, and Jun Kamei, Ryuji Yamaguchi, Yuichi Matsuda, Yoshio Sugii, Takashi Okauchi, Rie Yoshida, Maki Fujiwara, and Mayuko Nakano for their technical assistance. This study was supported by MEXT/JSPS KAKENHI Grant Numbers JP18K15353, JP21K07268, JP22H05521 (to KO), and, JP17H02219 (to TH), JP22H 05157 (to KI), JP19H05467 (to MT), and JP20H05955 (to TM), by JST PRESTO Grant Numbers JPMJPR22S3 (to KO), and JPMJPR2128 (to K Majima) and by AMED Grant Numbers JP23dm0307007 (to TH), JP21dm0107146 (to TM), JP20dm0307021 (to KI), and JP21dm0207077 (to MT). M.A.G.E and B.J.R were supported by the Intramural Research Program, NIMH/NIH/DHHS (project# ZIAMH 002032).

## Materials & Methods

### Subjects

Two male Japanese monkeys (Macaca fuscata) participated in the experiments (Mk #1: 7.2 kg; Mk #2: 6.8 kg; both aged 4 years at the beginning of experiments). The monkeys were kept in individual primate cages in an air-conditioned room. A standard diet, supplementary fruits/vegetables, and a tablet of vitamin C (200 mg) were provided daily. All experimental procedures involving the monkeys were carried out in accordance with the Guide for the Care and Use of Nonhuman primates in Neuroscience Research (The Japan Neuroscience Society; https://www.jnss.org/en/animal_primates) and were approved by the Animal Ethics Committee of the National Institutes for Quantum Science and Technology.

### Viral vector production

Mk #1 was co-injected with two AAV vectors, one expressing hM4Di and the other expressing GFP (AAV2–CMV–hM4Di and AAV2–CMV–AcGFP; 2.3 × 10*e*13 and 4.6 × 10*e*12 particles/mL, respectively). Mk #2 was injected with an AAV vector expressing both hM4Di and GFP (AAV2.1–CaMKII–hM4Di–IRES–AcGFP, 1.0×1013 particles/mL). AAV vectors were produced by a helper–free triple transfection procedure, and were purified b y a ffinity ch romatography (GE Healthcare, Chicago, USA). Viral titer was determined by quantitative PCR using Taq-Man technology (Life Technologies, Waltham, USA)(43).

### Surgical procedures and viral vector injections

Surgeries were performed under aseptic conditions in a fully equipped operating suite. We monitored body temperature, heart rate, SpO2, and tidal CO2 throughout all surgical procedures. Monkeys were immobilized by intramuscular (i.m.) injection of ketamine (5–10 mg/kg) and xylazine (0.2–0.5 mg/kg) and intubated with an endotracheal tube. Anesthesia was maintained with isoflurane (1%–3%, t o e ffect). B efore s urgery, m agnetic r esonance (MR) imaging (7 tesla 400 mm/SS system, NIRS/KOBELCO/Brucker) and X-ray computed tomography (CT) scans (Accuitomo170, J. MORITA CO., Kyoto, Japan) were performed under anesthesia (continuous intravenous infusion of propofol 0.2–0.6 mg/kg/min). Overlay MR and CT images were created using PMOD® image-analysis software (PMOD Technologies Ltd, Zurich, Switzerland) to estimate stereotaxic coordinates of target brain structures. After surgery, prophylactic antibiotics and analgesics (cefmetazole, 25–50 mg/kg; ketoprofen, 1–2 mg/kg) were administered.

The bilateral OFCs (BA11 & BA13) of each monkey were injected with the AAV vectors (Fig. 1A). The injections were performed under direct vision using the same types of surgical procedures as in a previous study(44). Briefly, after retracting skin, galea, and muscle, the frontal cortex was exposed by removing a bone flap and reflecting the dura mater. Then, handheld injections were made under visual guidance through an operating microscope (Leica M220, Leica Microsystems GmbH, Wetzlar, Germany), with care taken to place the beveled tip of a microsyringe (Model 1701RN, Hamilton) containing the viral vector at an angle oblique to the brain surface. The needle (26 Gauge, PT2) was inserted into the intended area of injection by one person and a second person pressed the plunge #1 and MK #2 via 53 and 49 tracks, respectively.

### PET imaging

PET imaging was conducted as previously reported(3). Briefly, PET scans were conducted before injection of vectors and at 45 days after injection for both monkeys. PET scans were performed using a microPET Focus 220 scanner (Siemens Medical Solutions USA, Malvern, USA). Monkeys were immobilized by ketamine (5–10 mg/kg) and xylazine (0.2–0.5 mg/kg) and then maintained under anesthetized condition with isoflurane (1%–3%) during all PET procedures. Transmission scans were performed for approximately 20 min with a Ge-68 source. Emission scans were acquired in 3D list mode with an energy window of 350–750 keV after intravenous bolus injection of [11C]clozapine (for Mk1; 375.5–394.7 MBq) or [11C]DCZ (for MK2; 324.9–382.3 MBq). Emission data acquisition lasted 90 min. To estimate the specific binding of [11C]DCZ in Mk2, regional binding potential relative to nondisplaceable radioligand (BPND) was calculated by PMOD® with an original multilinear reference tissue model (MRTMo). To visualize the expression of DREADDs, contrast (subtraction) of images taken before and 45 days after vector injection were created using PMOD for SUV (standardized uptake value) for Mk1 and BPND for Mk2 by investigating whether differential PET signals were observed at the target sites.

### Drug administration

DCZ (HY-42110; MedChemExpress) was dissolved in 2.5% dimethyl sulfoxide (DMSO, FUJIFILM Wako Pure Chemical Co.), aliquoted and stored at –30°C. For systemic i.m. injection, this stock solution was first diluted in saline to a final volume of 100 μg/kg. Fresh solution was prepared on each day of usage.

For microinfusion, DCZ was first dissolved in DMSO and then diluted in PBS to a final concentration of 100 nM. We prepared fresh solutions on the day of usage. We used two stainless steel infusion cannulae (outer diameter 300 μm; Muromachi-Kikai) inserted into each target region: rmCD and MDm, and ventral putamen for additional experiments (Fig. S7). Each cannula was connected to a 10-μL microsyringe (7105KH; Hamilton) via polyethylene tubing. These cannulae were advanced via guide tube by means of an oil-drive micromanipulator. DCZ solution or PBS was injected at a rate of 0.25 μL/min by auto-injector (Legato210; KD Scientific) for a total volume of 3 μL for each hemisphere. The injection volumes were determined based on a previous study reporting that injections of 3 μL and 1.5 μL resulted in a diameter of aqueous spread in the monkey brain of approximately 5–6 mm and 3–4 mm, respectively(45). We chose sufficient volumes to cover the hM4Di-positive terminal sites, which had diameters of 5–7 mm and 3–4 mm for the rmCD and MDm, respectively, as measured by increased PET signals. Because the MDm is located close to the midline, we placed the canulae laterally near the MDm so that the injected solution would diffuse into the entirety of the MDm (Fig. 3B). CT image was obtained to visualize the infusion cannulae in relation to the chambers and skull following each infusion. The CT image was overlaid on MR and PET images obtained beforehand using PMOD to verify that the infusion sites (tips of the infusion cannulae) were located in the target (presumed hM4Di-positive terminal regions identified as increased PET signals). The behavioral session began approximately 30 min after the end of the infusion and lasted approximately one hour. We performed at most one silencing experiment per week for one area.

### Behavioral tasks

The monkeys were tested with two versions of modified reversal learning tasks in which they were required to choose either of two visual stimuli (out of a set of five) presented on a computer screen. Behavioral testing was conducted in a sound-attenuated room. The monkeys sat on a monkey chair from which they could reach out one with hand to touch an LCD display placed in front of them. The behavioral task was controlled by a computer using commercially available software (Inquisit, Millisecond). A monkey initiated a trial by touching a sensor mounted on the chair, which caused a small white circle to appear in the center of the display. After a delay of 0.5 s, the circle disappeared and two stimuli of the five possible stimuli were presented simultaneously on the left and right side of the display. If the monkey touched either stimulus, it could receive a reward from the spout placed in front of its mouth. If the monkey released the touch lever before the presentation of visual stimuli, the trial was aborted and repeated after a 3-4 s inter-trial interval. Each stimulus was associated with 1, 2, 3, 4, or 5 drops of juice.

In the NOVEL task, a new set of visual stimuli was introduced each session, which required the monkeys to learn a new set of stimulus-reward associations. A daily session consisted of 90 acquisition-phase trials, followed by the reversal of the stimulus-reward associations, and then 210 post-reversal trials. The reversal was conducted such that a stimulus previously associated with 1 drop of juice became associated with 5 drops of juice, and one associated with 2 drops became associated with 4 drops, and vice versa. The combination of visual stimuli seen on each trial was pre-determined pseudorandomly so that each combination appeared once every 10 trials in a round-robin fashion.

In the FAMILIAR task, a fixed sets of five visual stimuli were used throughout the experiments. This ensured that the monkeys became familiar with all possible stimulus-reward associations for the two sets before and after reversals. If the optimal choice rate (i.e., the proportion of trials in which the option associated with the greater reward was chosen) in 30 consecutive trials passed 76%, the associations were reversed. Daily sessions comprised 300 and 400 trials for Mk1 and Mk2, respectively. Both monkeys were trained on the NOVEL task and then the FAMILIAR task. As a control, the monkeys were tested on a simple reversal learning task (Extended Data Fig. 5) in which only two novel stimuli were introduced in each session. The stimuli were associated with 5 drops of juice reward or no reward.

One monkey (Mk2) was also tested on a reinforcer devaluation task and on a modified version of the Wisconsin Card Sorting task, as previously described20,24. Briefly, in the reinforcer devaluation task, the monkey was required to choose one of two objects that were placed above two holes located in a wooden plate. For one set of objects, food 1 (peanut) was delivered when the object was selected, whereas for the other set of objects, food 2 (raisin) was delivered. The associations between the objects and reward type were fixed throughout training. The monkey was tested on 4 consecutive days in a week: review of object-reward associations (Day 1), baseline choice test (Day 2), review of object-reward associations (Day 3), and choice test following selective satiation (devaluation of a food) with vehicle or DCZ administration 15 min before devaluation (Day 4). The monkey’s ability to adaptively shift away from choosing objects associated with the devalued food was calculated as the “proportion shifted” as below, Proportion shifted=((F1N-F1D)+(F2N-F2D))/((F1N+F2N)) where F1 and F2 represent choices associated with the two food types (peanut and raisin) in each week in which that food type was devalued, and D and N respectively represent the data for devaluation (Day 4) and baseline (Day 2).

### Statistics

Throughout the manuscript, we compared learning rate and optimal choice rate for each monkey between conditions (vehicle and DCZ injections). Comparisons were analyzed with two-way repeated-measures ANOVA (treatment × monkey) to examine the effect of each treatment and individual differences. The analyses were conducted using GraphPad Prism 9. For the NOVEL task, the optimal choice rate was averaged across 10 trials of pseudo-random stimulus-reward combinations (see “Behavioral task”) for each session. In the FAMILIAR task, the optimal choice rate around the time of reversal was averaged across each reversal for each session. Reaction time in the NOVEL task was defined time between releasing the bar and touching the object on the screen, and data were averaged across 10 trials as described above. For statistical analysis of the post-reversal behavioral performance on the NOVEL task, we used the average optimal choice rate 100, 100, and 50 trials after the reversal for the OFC-, OFC-rmCD, and OFC-MDm silencing conditions, respectively. For the FAMILIAR task, we used the averages from the 10, 10, and 5 trials after the reversal, respectively. These trial numbers roughly corresponded to when behavioral deficits were observed if any.

### Histology and immunostaining

For histological inspection, monkeys were deeply anesthetized with an overdose of sodium pentobarbital (80 mg/kg, i.v.) and transcardially perfused with saline at 4°C, followed by 4% paraformaldehyde in 0.1 M phosphate buffered saline (PBS), pH 7.4. The brain was removed from the skull, postfixed in the same fresh fixative overnight, saturated with 30% sucrose in phosphate buffer (PB) at 4°C, and then cut serially into 50-μm-thick sections with a freezing microtome. For visualization of immunoreactive GFP signals (co-expressed with hM4Di), a series of every 6th section was immersed in 1% skim milk for 1 h at room temperature and incubated overnight at 4°C with rabbit anti-GFP monoclonal antibody (1:500, G10362, Thermo Fisher Scientific) i n PBS containing 0.1% Triton X-100 and 1% normal goat serum for 2 days at 4°C. The sections were then incubated in the same fresh medium containing biotinylated goat anti-rabbit IgG antibody (1:1,000; Jackson ImmunoResearch, West Grove, PA, USA) for 2 h at room temperature, followed by avidin-biotin-peroxidase complex (ABC Elite, Vector Laboratories, Burlingame, CA, USA) for 2 h at room temperature. For visualizing the antigen, the sections were reacted in 0.05 M Tris-HCl buffer (pH 7.6) containing 0.04% diaminobenzidine (DAB), 0.04% NiCl2, and 0.003% H2O2. The sections were mounted on gelatin-coated glass slides, air-dried, and cover-slipped. A portion of the other sections was Nissl-stained with 1% Cresyl violet. Images of sections were digitally captured using an optical microscope equipped with a high-grade charge-coupled device (CCD) camera (Biorevo, Keyence, Osaka, Japan).

### Model-fitting analysis

We constructed two mathematical models termed “EXP” and “INF” to investigate whether monkey behavior could be explained by experience- or inference-based strategies. In the EXP model, the probability that the learner (i.e., monkey) chooses stimulus Si when stimuli Si and Sj are presented (i,j=1,…,5) is given by

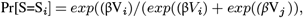

where β is a parameter controlling the exploration-exploitation trade-off (inverse temperature), and Vi is the subjective value of stimulus Si, which is interpreted as the amount of reward expected by the learner after choosing Si. Vi is updated after each trial based on the Rescorla–Wagner rule, which applies the stochastic gradient-descent algorithm to minimize the squared error between the expected reward for the chosen stimulus and the experienced reward. The update rule after choosing stimulus Si is given by

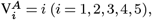

where α is the learning rate and *r*_*obs*_ is the amount of the reward provided in the current trial. All Vi (i=1,,5) were assumed to be zero at the beginning of each session, and the learning rate and inverse temperature were fitted to the behavioral data in each session via maximum likelihood estimation. In the INF model, we assume that the learner knows that the stimulus-reward association takes one of the two possible patterns. Taking this assumption, we prepared two sets of Vi:

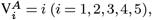

for pattern A, and

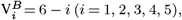

for pattern B. In the INF model, the learner’s choice is assumed to follow a mixture of the two softmax distributions corresponding to the two possible stimulus-reward association patterns. The probability that stimulus Si is chosen when stimuli Si and Sj are presented is given by

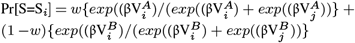

where β is the inverse temperature and w is a weight parameter adaptively learned based on the experienced rewards. If w = 1, the learner’s choice is assumed to follow the softmax distribution corresponding to stimulus-reward association pattern A, and if w = 0, the choice is assumed to follow the softmax distribution corresponding to stimulus-reward association pattern B. Thus, this weight parameter can be interpreted as a parameter representing the confidence that the current stimulus-reward association is pattern A out of the two possible patterns. Unlike in the EXP model, in the INF model, the inputs to the softmax function are fixed, and the parameter weighting the two softmax distributions is updated after each trial. Like the EXP model, the weight parameter is updated by the update rule derived from the stochastic gradient-descent algorithm, which minimizes the squared error between the expected reward for the chosen stimulus and the experienced reward. The update rule after choosing stimulus Si is given by

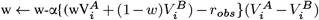

where α is the learning rate and robs is the observed reward in the current trial. w is assumed to be 0.5 at the beginning of each session and is assumed to take a value in the range of [0.0 1.0] at each update by replacing its value with 0.0/1.0 if it is smaller/larger than 0.0/1.0. As in the EXP model, the learning rate and inverse temperature were fitted to the behavioral data in each session via maximum likelihood estimation.

Because we were interested in the updating behavior during the post-reversal period, we modified the two models described above for the current study. In the modified models, the trials in each session were divided into two groups: 1) the trials in the post-reversal period, and 2) others. The post-reversal period consisted of all trials after the reversal in the NOVEL task, and the five trials after each reversal in the FAMILIAR task. While the learning rate in the basic models is fixed in each session, in the modified models we assumed that it took a different value each of the two trial groups. The learning rate for the post-reversal period is considered to better reflect the updating behavior, and a total of three parameters (two learning rates and inverse temperature) were fitted to the given behavioral data. Unless stated otherwise, the learning rates for the post-reversal periods derived from the modified models were reported in this paper.

To compare the ability of the EXP and INF models to explain the behavioral data, we performed Bayesian model comparison. Following the procedure in previous studies(46, 47), the Bayesian information criterion (BIC) was computed for each model and each session, then the Bayesian posterior probability for each model was computed based on those BIC values.

---

**Fig. S1.**
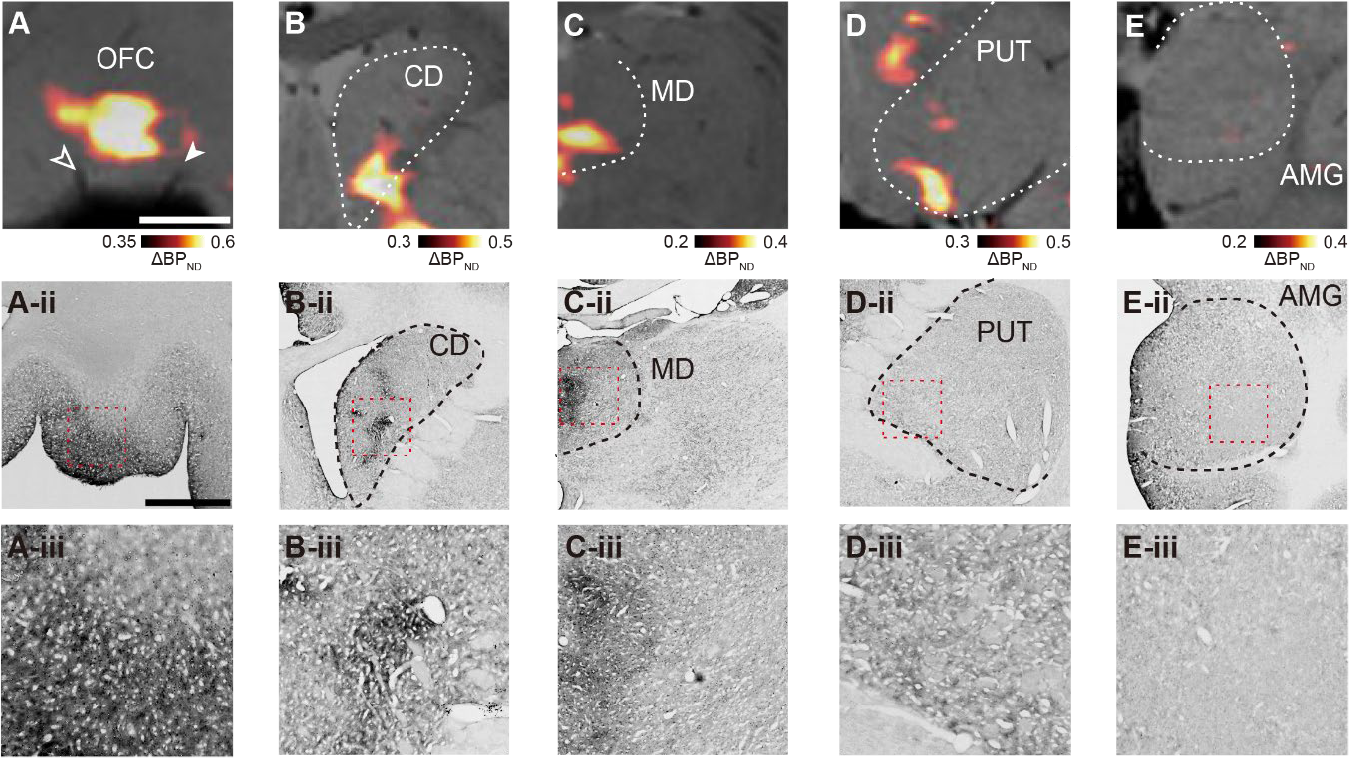
Expression of hM4Di in the OFC and its terminal sites. (**A** to **E**) *In vivo* visualization of hM4Di expression in the OFC (**A**), rmCD (**B**), MDm (**C**), putamen (**D**), and amygdala (**E**) obtained from Mk#2. Images are coronal PET contrasts showing specific binding of [^11^C]DCZ (contrast: after the introduction of hM4Di minus before the introduction), overlayed by MR images from Mk#2. The middle row visualizes corresponding DAB-stained sections showing immunoreactivity against a reporter protein (AcGFP), and the bottom row shows an enlarged view of the areas marked with red rectangles. Scale bars: 5 mm.

**Fig. S2.**
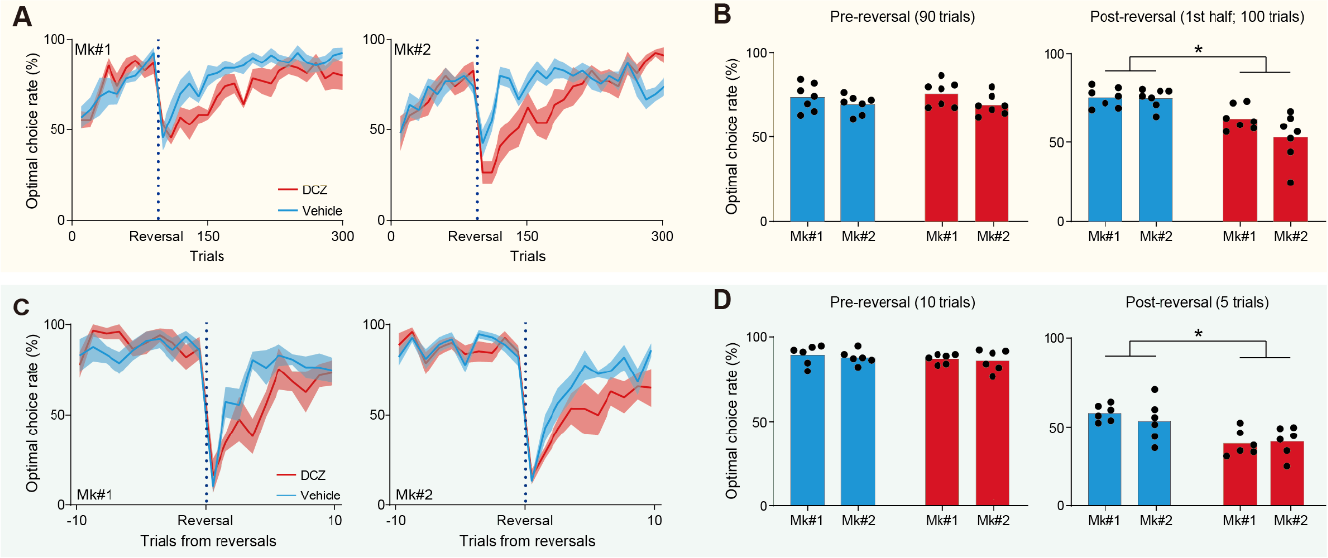
Effects of chemogenetic silencing of the bilateral OFC in each monkey. (**A**) Behavioral performance for Mk#1 (left) and Mk#2 (right) on the NOVEL task (N = 7 for each treatment). (**B**) Optimal choice rate for the pre-reversal phase (left, 90 trials; two-way ANOVA, treatment, *F*_(1,24)_ = 0.11, *p* = 0.74; subject, *F*_(1,24)_ = 4.4, *p* = 4.6 × 10^−2^; interaction, *F*_(1,24)_ = 0.21, *p* = 0.65) and the 1st half of the post-reversal phase (right, 100 trials, treatment, *F*_(1,24)_ = 28.0, *p* = 2.0 × 10^−5^; subject, *F*_(1,24)_ = 3.0, *p* = 0.10; interaction, *F*_(1,24)_ = 2.5, *p* = 0.12). (**C**) Behavioral performance for Mk#1 (left) and Mk#2 (right) on the FAMILIAR task (N = 6 for each treatment). (**D**) Optimal choice rate for the pre-reversal phase (left, 10 trials; two-way ANOVA, treatment, F_(1,20)_ = 0.36, p = 0.56; subject, *F*_(1,20)_ = 0.9, *p* = 0.34; interaction, *F*_(1,20)_ = 0.009, p = 0.93) and the post-reversal phase (right, 5 trials, treatment, *F*_(1,20)_ = 17.8, *p* = 4.2 × 10^−4^; subject, *F*_(1,20)_ = 0.25, *p* = 0.62; interaction, *F*_(1,20)_ = 0.68, *p* = 0.42). Asterisks: *p* < 0.05 for significant main effect of treatment.

**Fig. S3.**
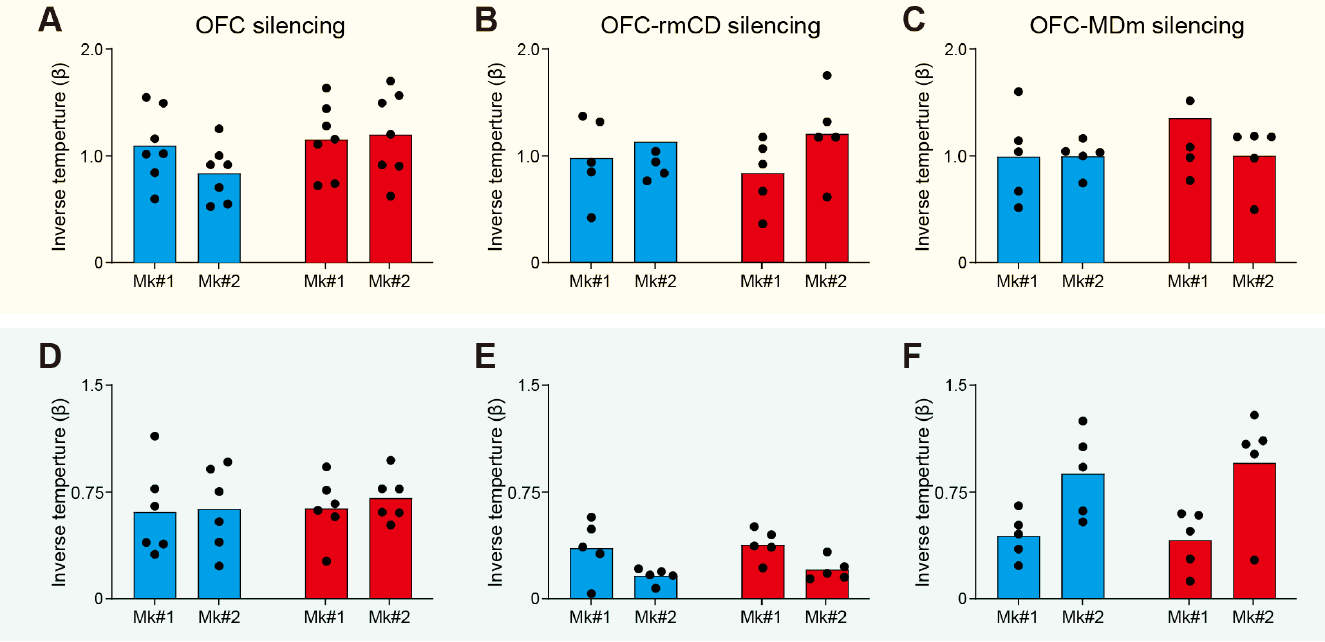
Estimated inverse temperature by model fitting analysis. (**A** to **C**) Inverse temperatures estimated by fitting performance on the NOVEL task to the EXP model after silencing the OFC (**A**) (two-way ANOVA, treatment, *F*_(1,24)_ = 2.7, *p* = 0.11; subject, *F*_(1,24)_ = 0.69, *p* = 0.41; interaction, *F*_(1,24)_ = 1.4, *p* = 0.24), the OFC-rmCD pathway (**B**) (treatment, *F*_(1,16)_ = 0.03, *p* = 0.86; subject, *F*_(1,16)_ = 1.9, *p* = 0.18; interaction, *F*_(1,16)_ = 0.33, *p* = 0.58), or the OFC-MDm pathway (**C**) (treatment, *F*_(1,16)_ = 0.94, *p* = 0.35; subject, *F*_(1,16)_ = 0.85, *p* = 0.37; interaction, *F*_(1,16)_ = 0.87, *p* = 0.37) for control vehicle (cyan) and DCZ treatment (red) in each monkey. Only data for the post-reversal phase are shown. (**D** to **F**) Inverse temperatures estimated by fitting to the performance on the FAMILIAR task to the INF model after silencing of the OFC (**D**) (two-way ANOVA, treatment, F_(1,20)_ = 0.21, p = 0.65; subject, *F*_(1,20)_ = 0.24, p = 0.63; interaction, *F*_(1,20)_ = 0.054, p = 0.82), the OFC-rmCD pathway (**E**) (treatment, *F*_(1,16)_ = 0.39, *p* = 0.54; subject, *F*_(1,16)_ = 11.0, *p* = 4.5 × 10^−3^; interaction, *F*_(1,16)_ = 0.026, *p* = 0.87), or the OFC-MDm pathway (**F**) (treatment, *F*_(1,16)_ = 0.034, *p* = 0.86; subject, *F*_(1,16)_ = 15.4, *p* = 1.2 × 10^−3^; interaction, *F*_(1,16)_ = 0.17, *p* = 0.68).

**Fig. S4.**
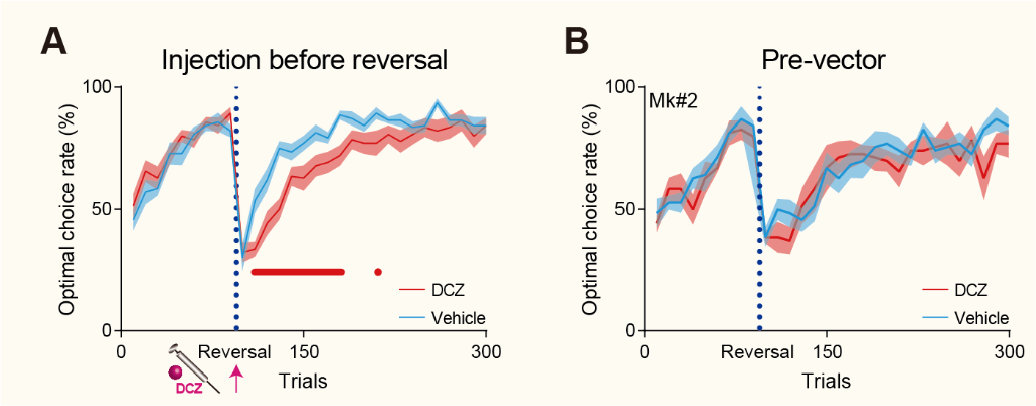
Effects of intramuscular DCZ administration just before the reversal and without hM4Di expression on the NOVEL task performance. (**A**) Behavioral performance on the NOVEL task (N = 7 for each treatment in each monkey) when DCZ was administered intramuscularly just before reversal. The task was interrupted for 5 min and DCZ was administered at the start of the 5-min break. (**B**) Behavioral performance on the NOVEL task (N = 7 for each treatment, Mk#2) before the introduction of hM4Di.

**Fig. S5.**
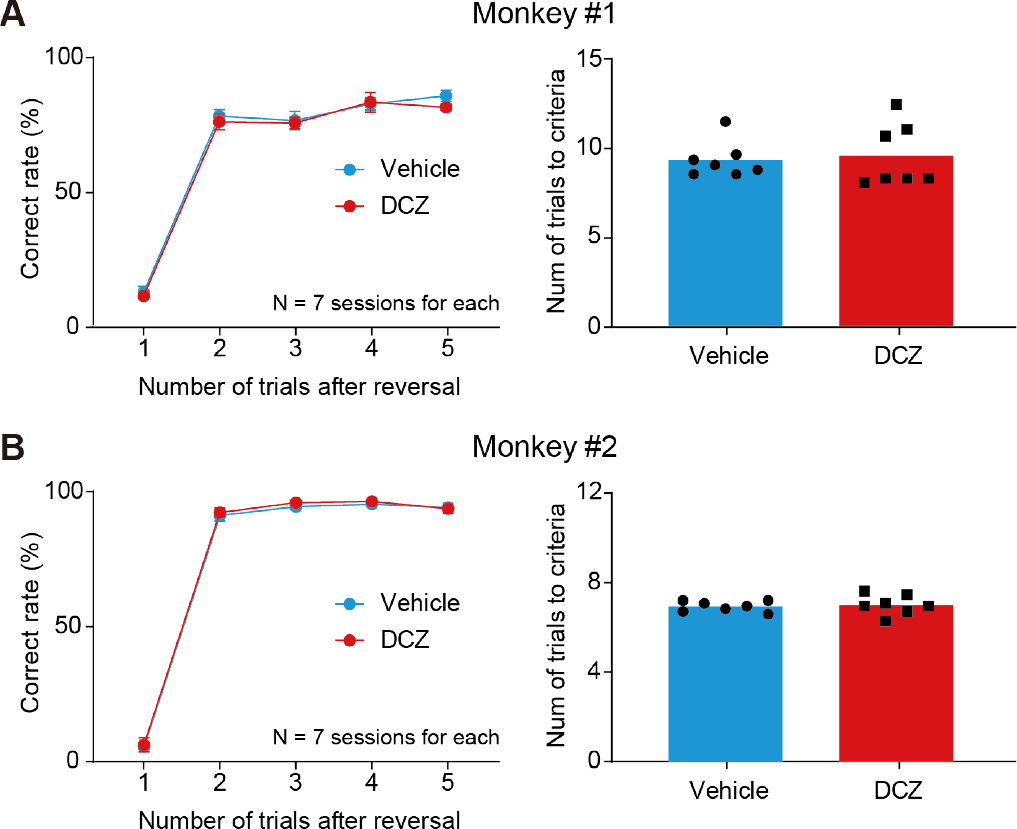
Effect of chemogenetically silencing the bilateral OFC on the two-arm reversal learning task. (**A** and **B**) Correct rate as a function of the number of trials after reversal (left) and the number of trials to reach criteria (right) for Mk#1 (**A**) and Mk#2 (**B**). There was no significant difference between vehicle and DCZ injection (Two-tailed Welch’s t-test, Mk#1, *t*_9.7_ = 0.33, *p* = 0.75; Mk#2, *t*_9.1_ = 0.37, *p* = 0.72).

**Fig. S6.**
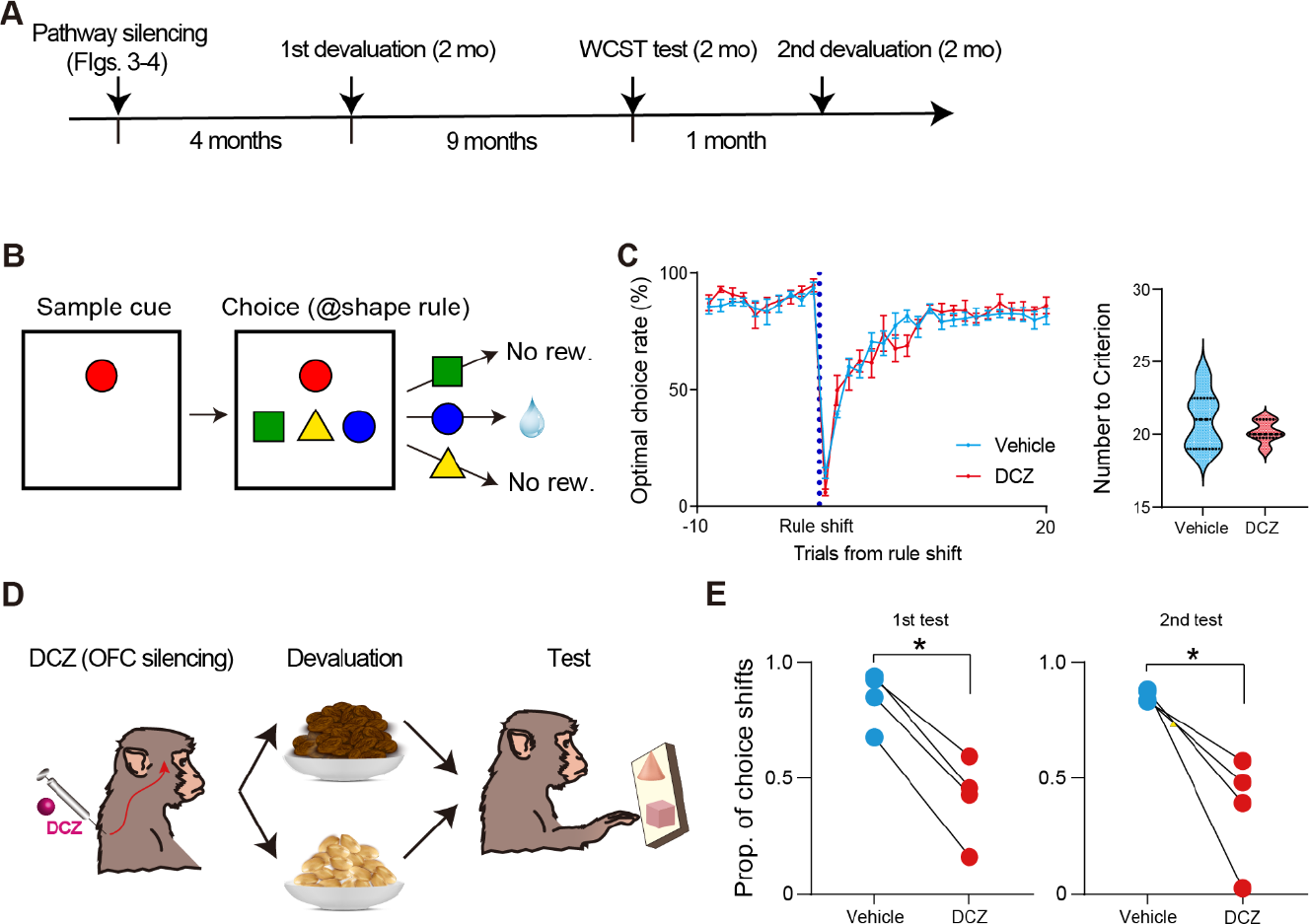
Effect of chemogenetically silencing the bilateral OFC on performance for the Wisconsin Card Sorting Test and the devaluation test. Schema of the Wisconsin Card Sorting Test. (**C**) Optimal choice rate (N = 6 for each treatment) as a function of trials for rule shift (left) and the number of trials to reach the criterion for rule shift (right). There was no significant difference in the number needed to reach criterion between vehicle and DCZ injection conditions (Two-tailed Welch’s t-test, *t*_6.4_ = 0.96, *p* = 0.37). The horizontal lines in each violin plot show the quartiles of the distributions. (**D**) Schema for the devaluation test. (**E**) Performance on the devaluation test for the 1st (left) and 2nd schedule (right), respectively (N = 4 for each treatment for both schedules). There was a significant difference in performance between vehicle and DCZ injections for both schedules (Two-tailed Welch’s t-test, 1st, *t*_3.1_ = 4.0, *p* = 2.7 × 10^−2^).

**Fig. S7.**
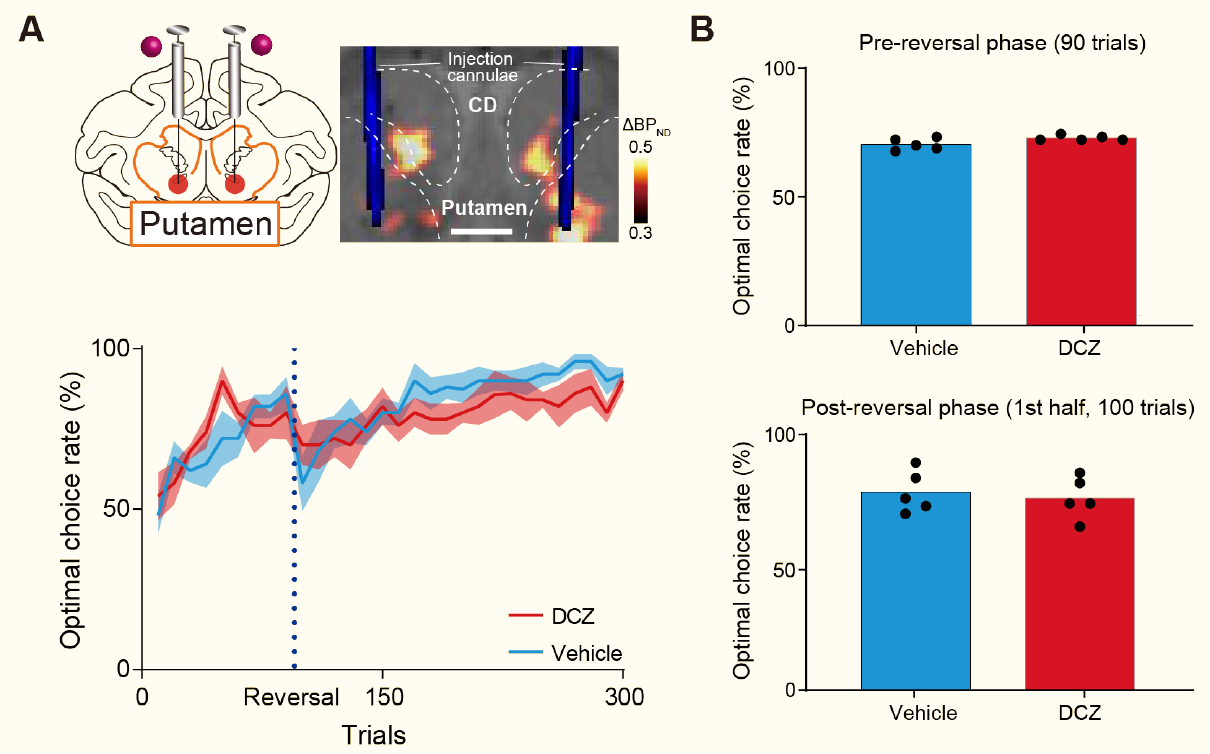
Effects of chemogenetically silencing the OFC-mPut pathway on NOVEL task performance. (**A**) Chemogenetically silencing the OFC-mPut pathway by local DCZ infusion into bilateral mPut, specifically at hM4Di-positive OFC terminals sites (top), and behavioral performance (bottom). N = 5 sessions for each treatment. Conventions are the same as in Fig. 2. (**B**) Averaged optimal choice rate in different task phases (Two-tailed Welch’s t-test, pre-reversal phase: *t*_5.4_ = 2.2, *p* = 0.08; 1st half of post-reversal phase: *t*_8.0_ = 0.46, *p* = 0.66).

**Fig. S8.**
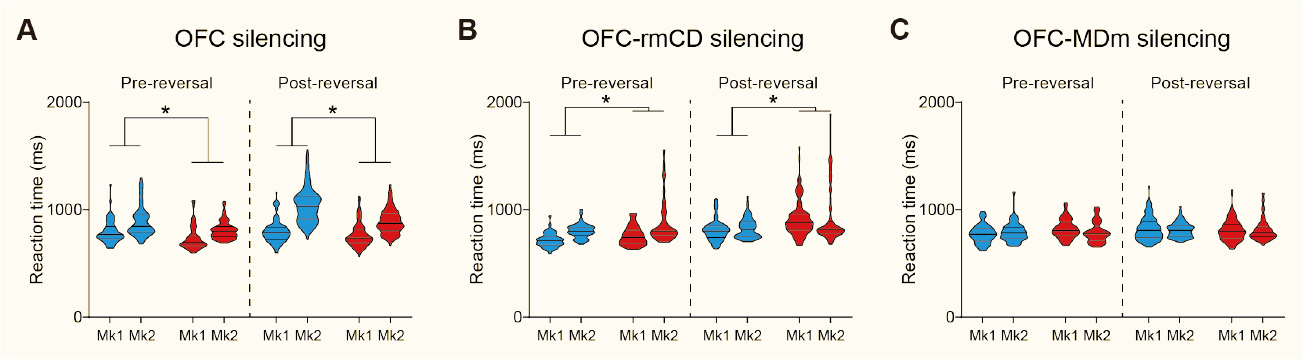
Effects of chemogenetically silencing the OFC, OFC-rmCD pathway, and OFC-MDm pathway on reaction time during the NOVEL task. (**A** to **C**) Reaction time in the pre-reversal phase (left) and post-reversal phase (right) after silencing the OFC (**A**), OFC-rmCD pathway (**B**), and OFC-MDm pathway (**C**) with DCZ treatment (red) or after control vehicle (cyan, no silencing) in each monkey. In both task phases, reaction times increased significantly after OFC silencing (pre-reversal phase: treatment, *F*_(1,248)_ = 25.3, *p* = 9.5 × 10^−7^; subject, *F*_(1,248)_ = 38.0, *p* = 2.8 × 10^−9^; interaction, *F*_(1,248)_ = 0.62, *p* = 0.43; post-reversal phase: treatment, *F*_(1,584)_ = 74.1, *p* = 6.8 × 10^−17^; subject, *F*_(1,584)_ = 325.7, *p* = 3.4 × 10^−58^; interaction, *F*_(1,584)_ = 74.1, *p* = 7.3 × 10^−10^), decrease significantly after silencing the OFC-rmCD pathway (pre-reversal phase: treatment, *F*_(1,176)_= 10.5, *p* = 1.5 × 10^−3^; subject, *F*_(1,176)_ = 32.6, *p* = 4.7 × 10^−8^; interaction, *F*_(1,176)_ = 0.77, *p* = 0.38; post-reversal phase: treatment, *F*_(1,416)_ = 37.8, *p* = 1.9 × 10^−9^; subject, *F*_(1,416)_ = 0.34, *p* = 0.56; interaction, *F*_(1,416)_ = 0.39, *p* = 0.53), and did not change after silencing the OFC-MDm pathway (pre-reversal phase: treatment, *F*_(1,176)_ = 2.7, *p* = 0.10; subject, *F*_(1,176)_ = 0.14, *p* = 0.71; interaction, *F*_(1,176)_ = 3.7, *p* = 0.06; post-reversal phase: treatment, *F*_(1,416)_ = 0.76, *p* = 0.38; subject, *F*_(1,416)_ = 0.22, *p* = 0.64; interaction, *F*_(1,416)_ = 0.58, p = 0.45). The horizontal lines in each violin plot show the quartiles of the distributions. Asterisks: *p* < 0.05 for significant main effect of treatment.

**Fig. S9.**
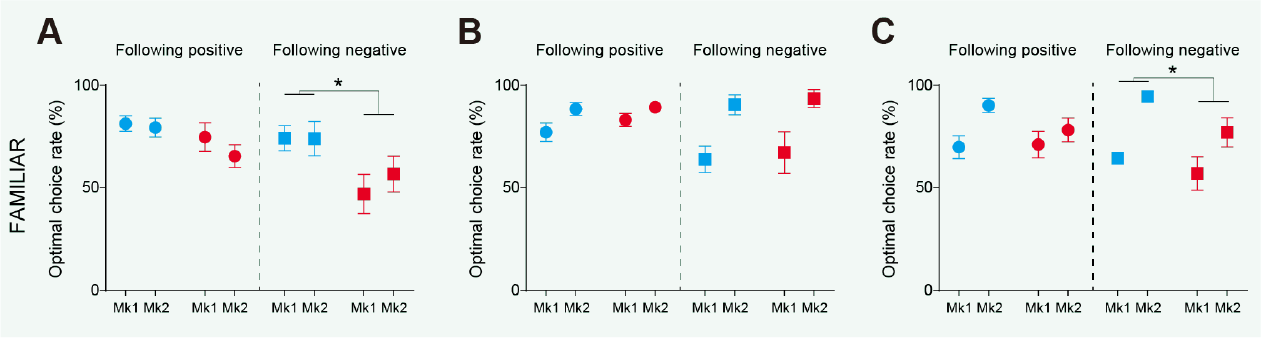
The effects of silencing the OFC, OFC-rmCD pathway, and OFC-MDm pathway on the sensitivity to past outcomes during the FAMILIAR task. (**A** to **C**) Averaged optimal choice rates for trials after the reversal in the FAMILIAR task following positive (left) and negative (right) outcomes after OFC silencing (**A**) (positive: treatment, *F*_(1,20)_ = 3.7, p = 0.069; subject, *F*_(1,20)_ = 1.1, *p* = 0.31; interaction, *F*_(1,20)_ = 0.47, *p* = 0.50; negative: treatment, *F*_(1,20)_ = 7.2, *p* = 1.4 × 10^−2^; subject, *F*_(1,20)_ = 0.33, *p* = 0.57; interaction, *F*_(1,20)_ = 0.36, *p* = 0.55), OFC-rmCD silencing (**B**) (positive: treatment, *F*_(1,16)_ = 1.0, *p* = 0.32; subject, *F*_(1,16)_ = 6.6, *p* = 2.1 × 10^−2^; interaction, *F*_(1,16)_ = 0.58, *p* = 0.46; negative: treatment, *F*_(1,16)_ = 0.21, *p* = 0.65; subject, *F*_(1,16)_ = 15.1, 1.3 × 10^−3^; interaction, *F*_(1,16)_ = 0.0004, *p* = 0.99), and OFC-MDm silencing (**C**) (positive: treatment, *F*_(1,16)_ = 0.40, *p* = 0.54; subject, *F*_(1,16)_ = 2.6, *p* = 0.13; interaction, *F*_(1,16)_ = 0.10, *p* = 0.76; negative: treatment, *F*_(1,16)_ = 6.0, *p* = 2.6 × 10^−2^; subject, *F*_(1,16)_ = 18.0, *p* = 6.0 × 10^−4^; interaction, *F*_(1,16)_ = 0.17, *p* = 0.68).

## Notes

### Competing Interest Statement

The authors have declared no competing interest.

